# Baseline Lateral Habenula Firing Rate Negatively Correlates with Increase in Ethanol Intake over Days in Adult Long Evans Rats

**DOI:** 10.1101/2021.01.27.428307

**Authors:** Shashank Tandon

## Abstract

While many adults consume alcohol, yet only some individuals are at a risk to develop alcohol use disorder (AUD). Variability in the risk for alcohol abuse is multifactorial and includes differences in behavioral and neuronal traits. The lateral habenula (LHb) has been shown to mediate aversive state-related behavioral responses. Interestingly, in both humans and rodents, depression-like symptoms are associated with high LHb activity. Additionally, there is a high co-morbidity between major depressive disorder and AUD. However, LHb lesions in rodents increase ethanol intake over time. Thus, we wanted to determine how baseline LHb activity correlates with ethanol intake over days. Specifically, we wanted to test whether individual variation in baseline LHb activity in ethanol-naive rats is related to home-cage ethanol drinking patterns. Hence, in this study, we determined the correlation between individual variability in baseline LHb neural activity, the negative-affective state-related ultrasonic vocalizations (USVs; 22-28 kHz), and the extent of ethanol intake over days. We first surgically implanted a unilateral 16-wire electrode array in the LHb of adult male Long Evans rats (n=11). Following recovery from surgery, rats were placed in sound-insulated chambers for two hours, where they were free to explore while we simultaneously recorded neuronal signals from their LHb and the USVs emitted by them. Next, these rats underwent an intermittent access to ethanol (IAE) paradigm, where they received 20% ethanol for 24 hours on alternate days (Monday-Wednesday-Friday) and ad-libitum water in their home cages for four weeks. The change in ethanol intake over days differed between rats, with some rats escalating their ethanol intake in the four weeks, while other rats showing no meaningful change in ethanol intake over days as compared to the first session. We found a significant negative correlation between average baseline LHb firing rates in rats and the changes in ethanol intake in the first week of IAE. Specifically, rats with higher baseline LHb firing rats, unlike rats with lower baseline firing rates, did not escalate their ethanol intake from the first session to the second and fourth ethanol sessions of the IAE paradigm. We also found a moderate positive correlation between the number of 22-28 kHz USVs and the average LHb firing rates of these rats. These results indicate that higher baseline LHb neuronal activity in normal adult ethanol naive rats is associated with decreased motivation to seek and consume ethanol during the early stages of ethanol consumption.

## Introduction

Alcohol abuse is a huge social health problem in the United States with around 5.8% or 14.4 million adults in the US ages 18 and older suffering from alcohol use disorder (AUD) in 2018 [1]. Interestingly, while many people consume alcohol, yet only some vulnerable individuals develop AUD while others do not [2]. The reasons for this difference in risk for alcohol abuse are multifactorial, including genetics, behavioral traits, and differences in metabolism [3, 4]. Likewise, individuals with certain neuronal traits are more at risk to develop AUD in the future [5]. Thus, understanding the neural basis for vulnerability to ethanol addiction is of paramount importance, as such insight will inform the development of preemptive strategies for individuals at high risk of AUD.

The lateral habenula (LHb) is a forebrain area implicated in aversion-mediated learning that has been suggested to play a role in modulating ethanol intake [6, 7]. The LHb receives inputs from limbic, basal ganglia and cortical areas and sends an excitatory projection to the rostromedial tegmental nucleus (RMTg), which in turn sends GABAergic inhibitory inputs to the dopaminergic neurons of the ventral tegmental area (VTA; [8–11]). LHb neurons also sends direct excitatory projections to dorsal raphe (DR) nucleus [12, 13]. Acting through the LHb–RMTg–VTA circuit, excitation of neurons in the LHb results in inhibition of VTA dopaminergic neurons [14–16]. Thus, LHb neuronal projections to the dopaminergic and serotonergic neurons can modulate motivation for any goal directed behavior.

LHb has been shown to regulate intake of drugs of abuse; *e.g.,* cocaine and nicotine [17–19]. Our prior studies found that LHb activity restricts escalation of ethanol intake. To elaborate, rats with lesions of the LHb consume more ethanol in an intermittent access to ethanol (IAE) paradigm and increase ethanol self-administration to a greater degree than do control rats [6]. In addition, ethanol-naïve rats with lesions of the LHb have attenuated ethanol-induced conditioned taste aversion (CTA) [7]. However, as LHb lesions in these studies were permanent, we could not delineate the relative contribution of baseline LHb activity (prior to ethanol use) and ethanol-induced LHb activity in regulating future ethanol consumption. Furthermore, our recordings of LHb activity in wild-type, ethanol-naive rats showed that high LHb neuronal activity decreases ethanol seeking as seen through decreased expression of ethanol-induced CTA [7, 20]. Thus, these rodent studies suggest that higher baseline LHb neuronal activity decreases ethanol-seeking behavior.

Interestingly, both human and rodent studies have shown LHb hyperactivity associated with depression like symptoms [9, 21, 22]. Additionally, a large percentage of people who suffer from depressive disorders also suffer from AUD [23–25]. Likewise, there is evidence for a correlation between acute negative emotional state and increased alcohol intake [26, 27].Thus, the alternate possibility is that high LHb activity is associated with high ethanol seeking and consumption.

Hence, it is important to delineate how baseline LHb activity correlates with future ethanol intake over time. The goal of this study, therefore, was to address this significant void in our knowledge by examining differences in LHb activity in individual rats prior to ethanol use and correlating it with ethanol use over days.

To measure the baseline affective state of the rats, we also simultaneously recorded the ultrasonic vocalizations (USVs) emitted by the rats along with the LHb neuronal activity. Rats emit 22-28 KHz frequency calls during distressed states, and these calls are initiated through the ascending cholinergic pathway [28, 29]. Rats also emit 50-55 KHz call during positive affective states, and these calls are initiated through the dopaminergic pathway [28, 29]. It is suggested that the mutual inhibitory interaction between the cholinergic and dopaminergic system initiates and maintains the affective states [28, 29]. USVs emitted by rats at these two different frequency bands have been used to determine the affective state of rats and predict the propensity of rats to increase their ethanol intake over time [30, 31]. We recorded USVs with LHb recordings to determine whether there is a correlation between baseline LHb neuronal activity and the number of negative-affect related USVs.

## Methods

Male Long-Evans rats (n=11; 300–350 g at the time of receipt; Charles-River, Wilmington, MA,) were used in the present experiments. Rats were single-housed in Plexiglas tub cages and maintained on a 12-hour light/dark cycle. Ad libitum access to food and water was available throughout all experimental procedures. All procedures used were approved by the University of Utah Animal Care and Use Committee and carried out in accordance with the National Institutes of Health Guide for the Care and Use of Laboratory Animals.

The timeline of the experiments were as follows: Rats following acclimatization first underwent surgery for electrode array implantation in the LHb. One week to ten days post-surgery they were placed in a sound-attenuated chamber for 2 hrs for USV and electrophysicology recording. The IAE paradigm started from the following Monday for four weeks. The rats were sacrificed after the twelveth ethanol session. If the array implant got detached from the skull prior to the 12^th^ ethanol session, the rat was immediately sacrificed and the ethanol intake in the last ethanol session was used for analysis.

The rats underwent surgery during which 16-wire electrode arrays was implanted in the LHb unilaterally (anteroposterior, −3.5 mm; mediolateral, 0.7 mm; ventral, −5.1 mm relative to bregma), as is routine in our laboratory [7]. Briefly, rats were anesthetized by isoflurane (5% and 2–3% for induction and maintenance, respectively). Craniotomy was performed, and the array was implanted stereotaxically and anchored to the skull with skull screws and dental cement. Following surgery, rats were administered buprenorphine (0.5 mg/kg) for long-term analgesia. Neo-Predef was applied locally to the surgical wounds. Seven days following recovery from surgery, we recorded neural signals from these rats for two hours while the rats were free to move in an operant chamber. Neuronal signals were recorded with a unity head stage amplifier, amplified 10,000-fold and captured digitally using commercial hardware and software (Plexon Instruments, Dallas, TX, USA). A single channel without discernible units was assigned as a reference electrode. Single units were identified by thresholding with a minimum signal-to-noise ratio of 3:1 and by using principal component analysis to resolve individual units by waveform shape, interspike interval and firing characteristics. The chamber had a USV microphone (CM16/CMPA, Avisoft Bioacoustics, Berlin, Germany) attached at 45 degrees angle to record USVs emitted by these rats in the two hours. The calls were recorded on Avisoft Recording Software (Avisoft Bioacoustics, Berlin, Germany).

Ethanol (Decon Labs, King of Prussia, PA) solutions were prepared in filtered tap water to a concentration of 20% (v/v) for use in the IAE paradigm. Voluntary ethanol consumption was measured using an IAE two-bottle choice paradigm [6]. Rats were given 24-hour access to ethanol and water in a two bottle choice paradigm in the home cage 3 times per week (Monday, Wednesday and Friday). Ethanol bottles were placed in home cages at noon on access days. Ethanol and water bottle positions within each cage were alternated in successive drinking sessions to minimize the effect of side preferences. On days in which ethanol was not presented, ad lib water was available. IAE was provided for a minimum of four weeks (12 drinking sessions). Ethanol and water intake were measured by weighing bottles before and after each 24-hour access period. These measures were used to calculate ethanol intake (normalized to weight, g/kg/24 h) and preference ([ethanol intake/total fluid intake]*100). Body weight and food intake were measured weekly during IAE. Following completion of the IAE paradigm, the rats were deeply anaesthetized with CO2 and the brains isolated. The brains were cryoprotected for sectioning and staining with cresyl violet to identify locations of electrode tips (pending).

Neurons signals obtained during recording were further sorted offline (Offline Sorter, Plexon Instruments). Analysis focused on baseline firing rate of the LHb neurons. The number of neurons recorded in each rat was similar (average = 14±1 neurons per animal). In all analyses, all neurons recorded in the LHb were used to determine the population average firing rates and to construct the corresponding population rate histograms. Analyses of neural firing were performed using Neuroexplorer (Nex Technologies, Madison, AL, USA) and Sigmaplot. For the USV recording analysis, the average number of counts/hr of the 22-28 KHz and 50-55 KHz frequency band for recorded USVs in each hour of the two-hour recordings was determined in all rats. The USVs were detected, analyzed, and counted using the DeepSqueak software [32]. We focused on the 22-28 kHz USVs and used 0.4 tonality threshold criteria to separate real USVs (> 0.4 tonality threshold) emitted by the rat from the background ultrasonic noise. The level of correlation between the baseline LHb firing rate and ethanol intake or change in ethanol intake was classified as very strong (0.80-1.0), strong (0.60-0.79), moderate (0.40-0.59), weak (0.20-0.39), and very weak (0.00-0.19) based on Pearson’s correlation (R) classification [33]. We chose the ethanol intake in first, second, fourth and eight session to measure correlation between baseline LHb firing rates and changes in ethanol intake over days.

## Results

With regard to ethanol intake during the IAE paradigm, we found that the change in ethanol intake over days differed between rats with some rats escalating their ethanol intake (Fig. 1A) while other rats showing no meaningful change in ethanol intake over the four weeks (Fig. 1B). Specifically, in seven rats, the 24 hr-ethanol intake for 20% ethanol escalated over days in the IAE paradigm from 4.4±0.8 g/kg/24 hrs in the first session to 7.6±0.5 g/kg/24 hrs in their last ethanol session (Fig. 1A). In the other four rats, the intake was 3.4±0.8 g/kg/24 hrs in the first session to 3.1±0.9 g/kg/24 hrs in their last ethanol session (Fig. 1A). Likewise, the change in ethanol preference of rats was similar; with seven rats showing increase in ethanol preference from 33+9% in the first session to 66±9% in their last ethanol session (Fig. 1C), with no meaningful change in four rats from the first to their last ethanol session (first session, 41±14%; last ethanol session, 31±10%,; Fig. 1D).

**Fig 1.**
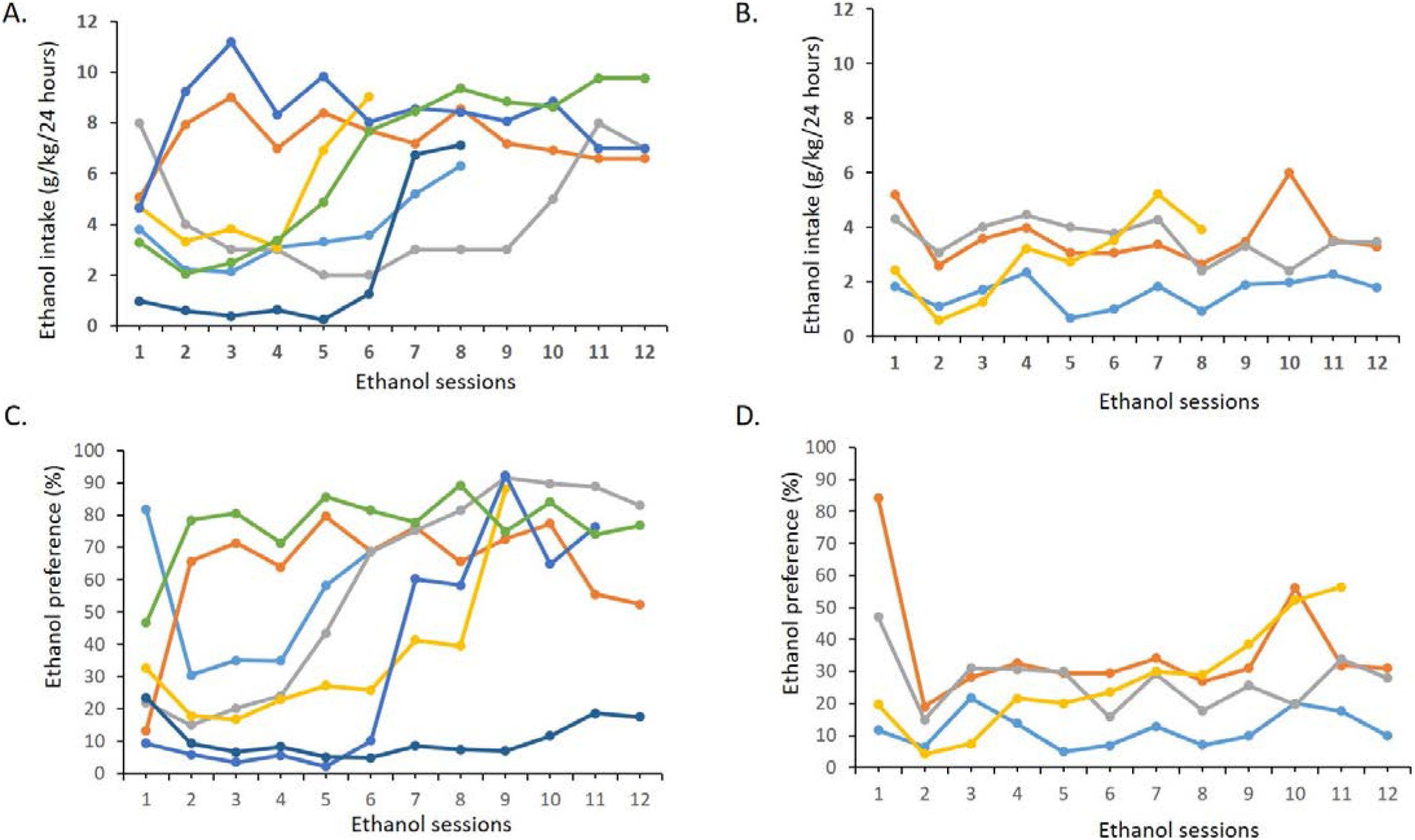
Voluntary ethanol consumption by individual rats in different sessions during the four-week intermittent access to ethanol paradigm. A) Rats in which ethanol consumption increased markedly over the course of 2-bottle choice drinking sessions (3 sessions/week for 4 weeks, 12 sessions total); B) Rats in which ethanol intake did not increase over the course of 2-bottle choice drinking sessions; C) Rats in which ethanol preference increased over the course of 2-bottle choice drinking sessions; D) Rats in which ethanol preference did not increase over the course of 2-bottle choice drinking sessions.

The change in ethanol intake and ethanol preference in rats from first to second session of IAE showed a very strong and strong negative correlation (ethanol intake, R = −0.85, P = 0.001; ethanol preference, R = −0.79, P = 0.009) with the baseline LHb firing rate, respectively (Fig. 2A and 2B). Specifically, rats with higher baseline LHb firing, unlike rats with lower baseline LHb firing rate, did not increase their ethanol intake and ethanol preference in the second session as compared to the first session (Fig. 2A and 2B).

**Fig 2.**
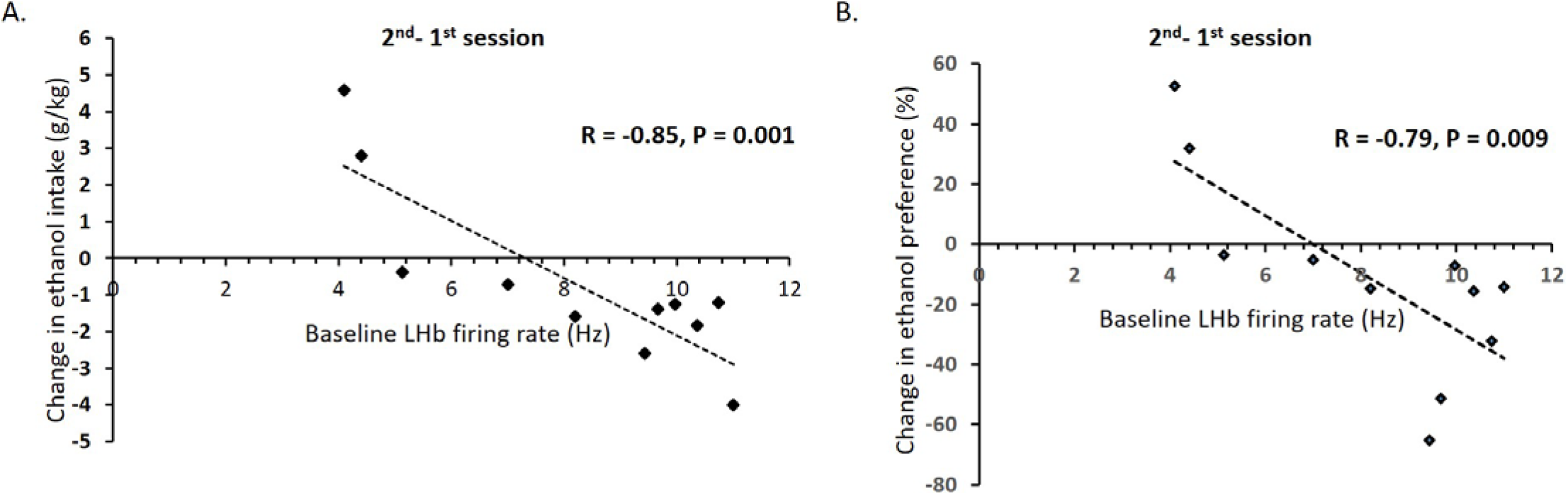
The correlation between baseline lateral habenula activity and change in ethanol consumption (A) and ethanol preference (B) for individual rats from the first to the second 2-bottle choice drinking session.

The change in ethanol intake in rats from first to fourth and first to eight session showed a strong (R = −0.66, P = 0.02) and moderate negative correlation (R = −0.57, P = 0.08) with the baseline LHb firing rate, respectively (Fig. 3A and 3B). Specifically, the increase in ethanol intake in rats with higher LHb firing rate was less compared with those with lower baseline LHb firing rate in the later sessions of the IAE paradigm.

**Fig 3.**
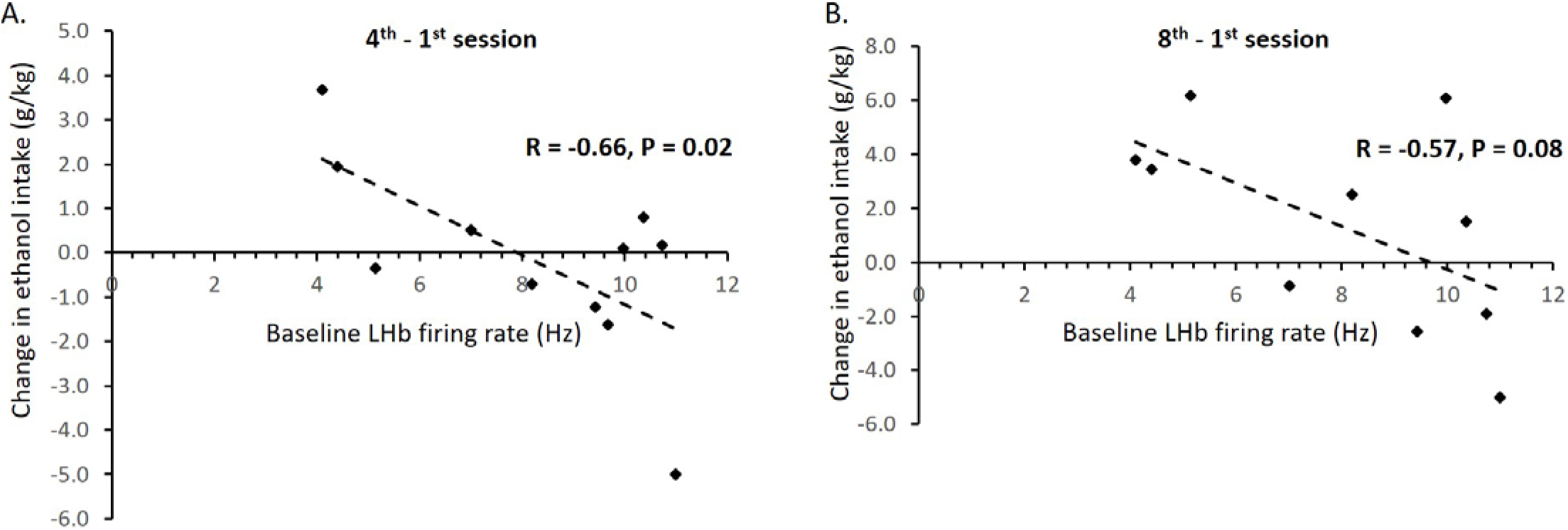
The correlation between baseline lateral habenula neuronal activity and change in ethanol consumption for individual rats from the first to fourth (A) and first to eight (B) 2-bottle choice drinking session.

When we compared ethanol intake in each session with baseline LHb firing, we found that the ethanol intake by rats in the second, fourth and eight session showed a moderate negative correlation (second session, R = −0.52, P = 0.1; fourth session, R = −0.41, P = 0.21; eight session, R = −0.52, P = 0.09) with the baseline LHb firing rate (Fig. 4B, C and D). However, the ethanol intake in the first session showed a very weak positive correlation (R = 0.03, P = 0.36) with the baseline LHb firing rate (Fig. 4A). These results show that baseline LHb activity does not modulate the first time ethanol drinking but likely negatively influences ethanol drinking in the later sessions.

**Fig 4.**
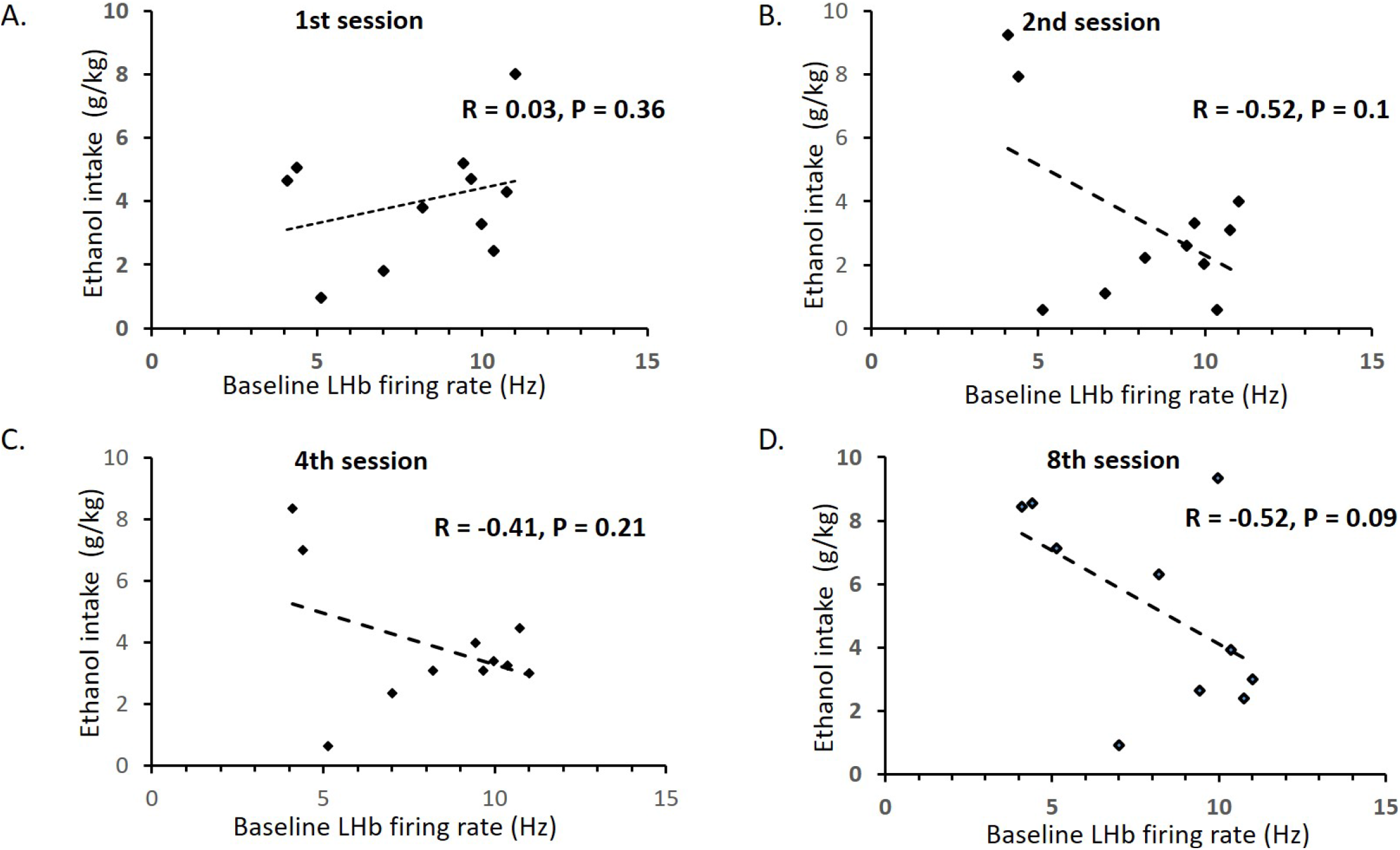
The correlation between baseline lateral habenula activity and ethanol consumption in the first (A), second (B), fourth (C) and eight session (D) of the IAE paradigm.

We also found that there is a moderate correlation (R = 0.55, P = 0.19) between baseline LHb activity and simultaneously recorded negative-affect associated 22-28 kHz USVs (Fig 5).

**Fig 5.**
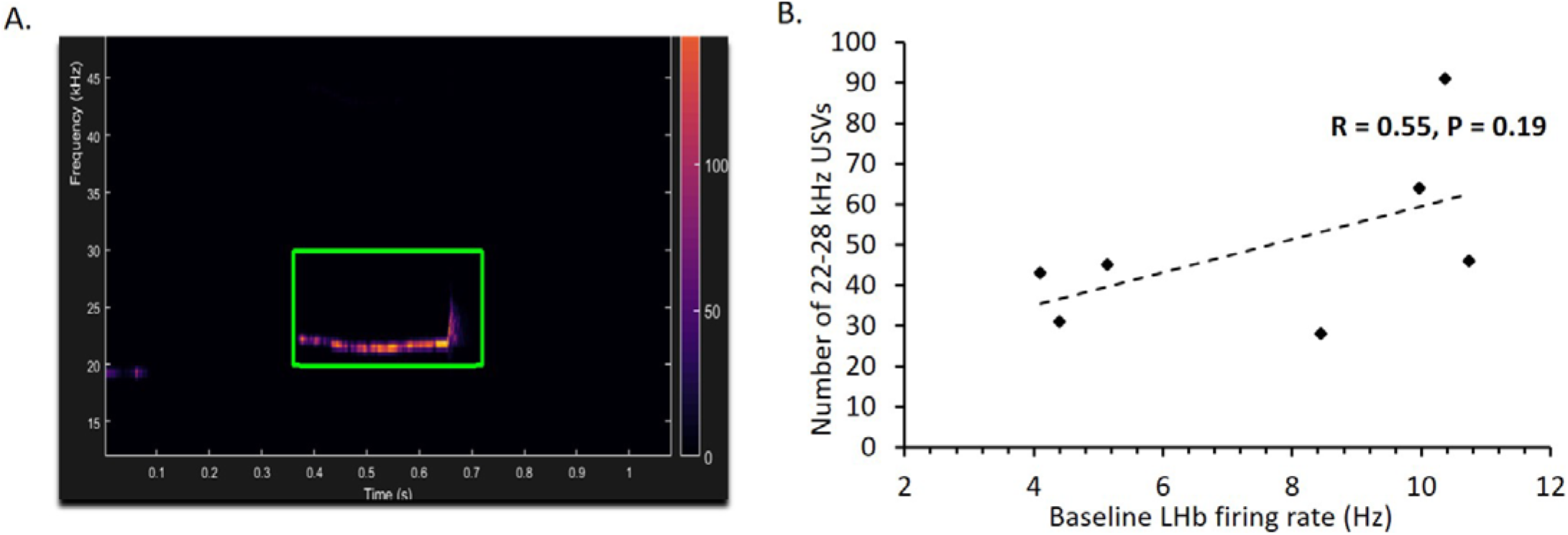
The correlation between the baseline lateral habenula firing and the number of 22-28 kHz calls. A) A USV call at 22 KHz isolated using the DeepSqueak software; B) Correlation between number of 20-22 KHz and Lateral habenula firing in the 2 hr recording.

## Discussion

These results are the first to show a correlation between individual variability in baseline LHb activity in normal adult rats to rate of increase in ethanol consumption over days. This indicates that baseline levels of LHb activity likely plays a modulatory role in the motivation to seek and consume ethanol during early stages of ethanol drinking. While we found that the correlation between baseline LHb activity and ethanol intake over later sessions is of a lesser degree, the trend was in the same direction as the early sessions - rats with lower baseline firing rate drank more ethanol in later sessions as compared with rats with higher baseline LHb firing rate. Interestingly, there was no meaningful correlation between baseline LHb firing rate and the first ethanol session, indicating that baseline LHb firing rate is not related to motivation to drink ethanol (a novel reward) in ethanol naïve rats.

As depression, associated with high LHb activity [9, 21, 22], leads to self-medication on ethanol [23, 24], we anticipated that higher LHb activity in rats would correlate with more ethanol drinking over days. Likewise, the negative symptoms e.g. anxiety and anhedonia during ethanol withdrawal are associated with a decrease in VTA dopaminergic activity [34–38], and an increase in RMTg neuronal activity [39] and DR neuronal activity [40], which suggests higher activity in the LHb neurons during withdrawal. Thus, in late stages of ethanol use, high LHb activity is associated with increased ethanol seeking and consumption. However, based on the results of this study and our previous findings [6, 7], it seems likely that during early stages of ethanol drinking in normal adult rats, high baseline LHb activity is related to degree of aversive response to ethanol.

We found that the negative correlation is the strongest between baseline LHb firing rate and the change in ethanol intake from the first to second and fourth ethanol session as compared with the correlation between baseline LHb firing rate and the absolute value of ethanol intake in the second and fourth session. This indicates that baseline LHb firing rate in ethanol naïve rats likely modulates the response to ethanol intake in the first ethanol session. Ethanol naïve rats have a strong delayed aversive response to ethanol [7], along with the initial pharmacological rewarding effect of ethanol on dopamine release [41, 42], with individual variability in this aversive response [7]. The strong correlation between baseline LHb firing rate and the difference in ethanol intake in second vs. first session could be because this degree of aversive response to ethanol depends, in part, on the baseline levels of LHb activity. Specifically, rats with higher baseline LHb firing rate could have stronger aversive response to ethanol compared with rats with weak aversive response to ethanol. As activation of the LHb-RMTg-dopamine pathway causes acute behavioral avoidance [43, 44] and aversive conditioning [44–47], higher LHb neuronal activity may lead to avoidance of ethanol seeking.

This is the first time USV recordings have been combined with neural recordings as a tool to determine the relationship between the affective state of the rats and the neuronal activity from any brain area at any given time. Future studies need to be designed to record LHb firing rates and the USVs emitted by the rats prior to each 24-hr ethanol access period in the IAE paradigm to understand how LHb activity and affective changes over time with chronic ethanol intake. It needs to be mentioned that our study was focused on individual differences in outbred rats with no other prior stress experience; and the next step would be to determine how baseline LHb neuronal activity and ethanol intake differs in rat with depression-like symptoms that have undergone stress [48–50] compared with normal rats.

## Acknowledgement

This work was supported by 1R03AA026758-01A1 grant to Shashank Tandon from the National Institute on Alcohol Abuse and Alcoholism (NIAAA). I would also like to acknowledge Dr. Kristen Keefe and members of her lab for their support during this project.

